# Size-based ecological interactions drive food web responses to climate warming

**DOI:** 10.1101/430082

**Authors:** Max Lindmark, Jan Ohlberger, Magnus Huss, Anna Gårdmark

## Abstract

Predicting the impacts of climate change on animal populations and communities requires understanding of feedbacks between direct physiological responses and indirect effects via ecological interactions. Food-dependent body growth and within-species size variation have major effects on dynamics of populations and communities through feedbacks between individual performance and population size structure. Moreover, evidence suggests a link between temperature and population size structure, but we lack an understanding of how this is mediated by species interactions when life history processes are food-dependent. Here, we use a dynamic stage-structured biomass model with food-, size- and temperature-dependent life history processes to assess how temperature affects coexistence, stability and size structure in a tri-trophic food chain. We show that predator biomass densities decline with warming either gradually or in the form of collapses, depending on which consumer life stage they predominantly feed on. Collapses occur when warming destabilizes the community and induces alternative stable states via Allee effects, which emerge when predators promote their own food source through predation. By contrast, warming at low temperatures stabilizes the community as limit cycles turn to fixed point dynamics, unless predators feed only on juveniles. Elevated costs of being large in warmer environments accelerate the decline in predator persistence and mean body size of the community. These results suggest that predator persistence in warmer climates may be lower than previously acknowledged when accounting for size- and food-dependence of life history processes, and that interactions within and between species can mediate the effects of warming on food web stability.

**Significance:** Climate warming is altering the dynamics and structure of aquatic ecosystems worldwide. Predicting food web reorganization under rising temperatures requires an understanding of physiological responses and ecological interactions of organisms, both of which depend on body size. We show that size variation within species, food-dependent growth and ecological interactions critically affect how food chains respond to warming. Specifically, warming can stabilize or destabilize food chains and expose predators to increased risk of sudden collapses, resulting in alternative stable food web states. Increasing temperatures can cause abrupt reductions in mean community body size, primarily due to loss of top predators. The potential loss of biodiversity and shifts in ecosystem stability are among the major challenges caused by a warming climate.

## Introduction

Predicting the impacts of climate change on natural food webs requires mechanistic understanding of organisms’ physiological responses to warming and how these translate to the population and community level. An individual’s metabolism, and related ecological traits including feeding, mortality and population growth rate, depend strongly on body size and temperature (1–3). Mechanistic models based on metabolic scaling theory have increased our understanding of how warming affects populations and communities in terms of (i) community size structure (4–6), (ii) strength of trophic interactions (7–9), (iii) food chain length (8, 10) and (iv) stability (10–12). The effects of temperature on (ii)-(iv) can largely be predicted from the relative temperature sensitivity of biomass gains (feeding) and losses (metabolism, mortality) – hereafter referred to as energetic efficiency – and resource productivity (8, 10–14). Specifically, increased energetic efficiency with temperature is generally predicted to have a destabilizing effect on communities and decreased efficiency a stabilizing effect (11). In addition, the latter scenario generally leads to predator extinction from starvation (10, 11). However, while these insights stem from mechanistic models that are typically based on body size dependence of individual-level processes, two fundamental aspects of body size are commonly overlooked when modeling effects of warming on populations and communities. First, the combination of within-species size variation and food-dependent life history processes (e.g. growth, development and reproduction) generates feedbacks between size structure and individual performance, ultimately affecting community dynamics (15). Second, the effects of warming differ between individuals, depending on life history stage and body size (16, 17). Therefore, we need to account for variation in body size within and between species to better understand the potential effects of warming on food web structure and stability (18).

Within-species size variation is not only universal in natural systems, but has major implications for the stability and structure of populations and communities because it leads to asymmetric competition between individuals of different sizes (ontogenetic asymmetry) (15). Ontogenetic asymmetry can lead to phenomena such as biomass overcompensation, which refers to an increase in standing stock biomass with mortality, and this is often life stage specific (19). Biomass overcompensation occurs when mortality releases a life stage from high density dependence, resulting in higher biomass production (greater than lost through that mortality). This phenomenon has been identified empirically in several studies (20–24). In the case of predation mortality, predators can thus cultivate biomass density of their own food source by inducing overcompensation in the prey, which can lead to an emergent Allee effect and alternative stable states when predator persistence relies on prey overcompensation (25, 26). Emergent Allee effects refer to a positive relationship between per capita predator population growth rate and their population density (Allee effect) that emerges from individual-level assumptions, such as size-scaling of feeding rates and maintenance costs, instead of predefined population dynamics. As a consequence, predator populations may be exposed to risks of sudden collapses from e.g. fishing mortality, from which they may not recover (26). Emergent Allee effects via food-dependent body growth have been demonstrated in a natural whole-lake experiment, where an overharvested predator population (brown trout, *Salmo trutta*, L.) could not control the size distribution of its then stunted prey (Arctic charr, *Salvelinus alpinus*, L.). However, culling of the stunted prey led to increased juvenile prey abundance on which predators fed, which shifted the community to a state with abundant predators in which predation kept the prey from a stunted state (22). The same mechanism has also been proposed to explain the lack of recovery of overfished Atlantic cod (*Gadus morhua*, L.) stocks, despite reduced fishing pressures (26–28). As food- and size-dependent body growth and within-species size variation are important for understanding dynamics of ecological communities, they are also key factors determining the effects of climate change on food webs.

In addition to affecting community structure and dynamics, the effects of warming on individuals tend to be size-dependent, and this interactive effect is possibly stronger in aquatic compared to terrestrial systems (17, 29–31). Lines of evidence that support size-dependent temperature effects include directional changes in size composition towards smaller, energetically more efficient individuals (6, 32, 33), the across taxa observation that size at maturity declines with warming (temperature-size rule, TSR) (29, 34, 35), and that optimum temperatures for growth decrease with size (36, 37). Despite the observational evidence of various temperature-size interactions, only recently have the dynamical consequences of such interactions in warming environments been explored (31, 38–40). Theoretical studies suggest that community persistence (in terms of food chain length) generally increases if body sizes decrease in warmer environments, due to weakened interaction strengths between species (40). Smaller mean body sizes can also increase stability (return times) and thus resilience of consumer-resource systems, which could buffer against extinctions (39). However, it is not known whether the effect of temperature-size interactions on stability and persistence holds when the change in size (or performance of a given size or life stage) with temperature depends on feedbacks between direct temperature effects, food availability and population size structure. This has not been studied for more than two interacting species (31, 38). As feedbacks between the biotic environment, individuals and population shape community size structure and dynamics, this knowledge gap limits our ability to predict how climate change impacts food chains and food webs (15, 18, 26, 31, 38).

Here we show how within- and between-species interactions mediate the direct physiological effects of warming on the stability and coexistence of a tri-trophic food chain, using a dynamic stage-structured biomass model with temperature-dependent vital rates. Our analyses generated novel predictions on community responses to changing temperatures that are due to food-dependent life history processes and species interactions; (i) whether warming stabilizes or destabilizes communities depends on the size preference of the predator, size structure in the consumer and the current temperature; (ii) warming can cause non-gradual declines (collapses) in predator populations due to Allee effects, which also induce alternative stable states in which predators either coexist with their prey or go extinct; (iii) increased energetic costs of being large in a warmer environment reduce the scope for predator persistence and the average community body size. These previously unrecognized temperature responses highlight that food-dependent life history processes and species interactions mediate the direct effects of warming on the dynamics and structure of ecological communities.

## Results

### Stabilizing and destabilizing effects of warming

Whether increasing temperatures stabilize or destabilize the food chain depends on the form of species interactions and the current temperature regime (Fig. 1–2). Warming at low temperatures have a stabilizing effect on the community as cyclic dynamics (“inverse enrichment cycles”) switch to fixed point dynamics (Hopf bifurcation at 12 °C in Fig. 1A-D) (Fig. 2) – unless the predator feeds more or less exclusively on juvenile consumers (Fig. 1E-H; Fig. 2) (*p* > 0.98). In contrast, at higher temperatures, e.g. reference temperature (19 °C), warming induces alternative stable states (bistability), with or without predators, when predators feed mainly on juveniles (Fig.1–2). In this case, warming has a destabilizing effect on community dynamics, as equilibrium dynamics switch from fixed point to bistability. By contrast, warming has no effect on the dynamical stability when predators feed more equally on both life stages. This means there is a region of juvenile feeding preference (0.72 < *p* < 0.98) where increasing temperatures initially stabilize the community, but induce bistability with additional warming (Fig. 2), i.e. the stability-temperature relationship depends on the current temperature regime. Irrespective of the predator’s feeding preference, the predator population declines in biomass density with warming until starvation (when predators go extinct). This decline can be gradual or in the form of a collapse, depending on which consumer life stage the predator feeds more on. Predators can thus experience sudden collapses when falling below a certain biomass threshold (Fig. 1H). The interaction between predators and consumers also regulates at which temperature predators go extinct, such that the extinction temperature decreases with increased feeding on juveniles (Fig. 2; SI Appendix, Fig. S9, Table S3-S4).

**Fig. 1.**
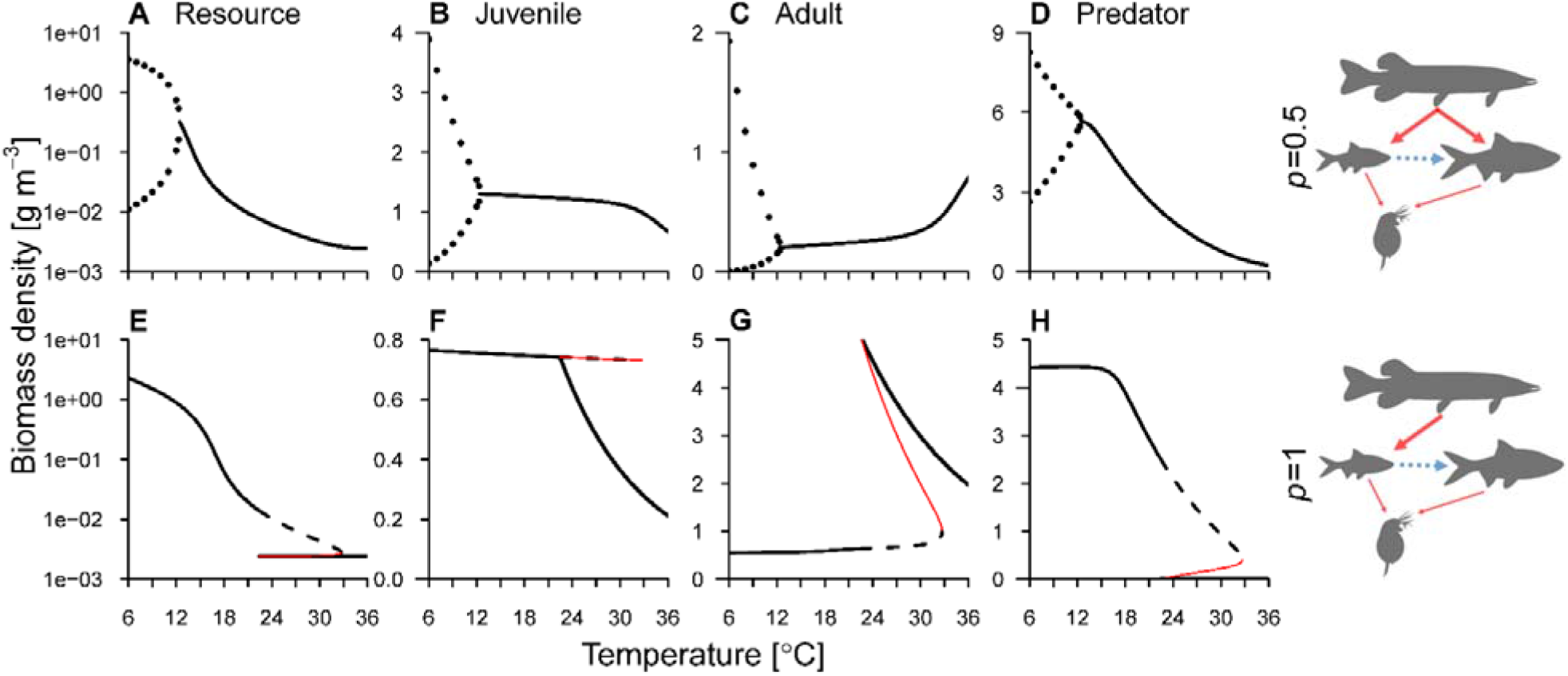
Effects of warming on food chain stability depend on ecological interactions. Equilibrium biomass densities of the resource (A, E), consumer life stages (B-C, F-G) and predator (D, H) as a function of temperature, given a predator feeding with equal intensity on both life stages (A-D) (*p* = 0.5) or exclusively on juveniles (E-H) (*p* = 1). Black lines (full and dashed) are stable equilibria and red thin lines are unstable equilibria (connecting the two stable branches in the bistable region), which separate the two stable equilibria when there are alternative stable states. Maximum and minimum biomass density of a stable limit cycle is shown with points (top row below ∼12 °C). Alternative stable states, where predators are either extinct or abundant, occur between ∼22-33 °C in E-H. Note the different scales on the y-axes and the logarithmic y-axis for resources densities. *E_R_max__* = −0.43, all other parameters have default values (SI Appendix, Table S1).

**Fig. 2.**
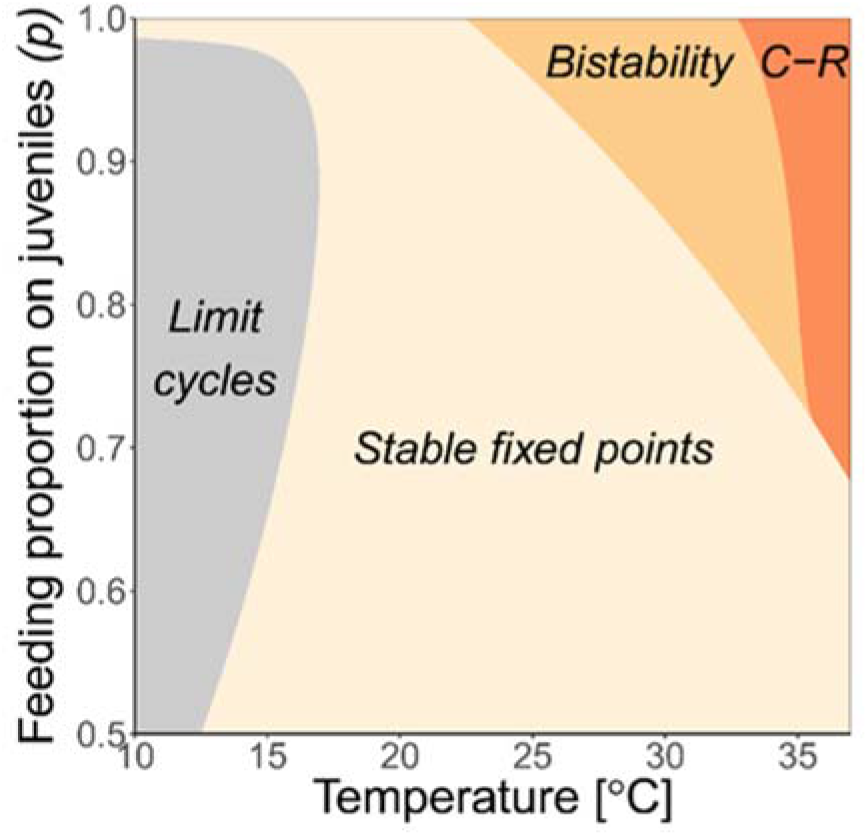
Community structure shifts with temperature and the predators’ feeding preference. In gray regions all species in the food-chain exhibit stable population cycles, off white corresponds to stable predator-consumer-resource states, beige shows bistable regions where the food-chain exhibits alternative stable states with predators being either extinct or absent (here the lower temperature boundary of the region corresponds to the invasion boundary and the upper is the persistence boundary), and orange is the stable consumer-resource system where predators cannot persist.*E_R_max__* = −0.43, all other parameters have default values (SI Appendix, Table S1).

The alternative stable states at warmer temperatures are due to emergent Allee effects in the predator population, i.e. there is a positive relationship between predator biomass density and their individual performance. The mechanism is that when predators feed primarily on juveniles (*p* > 0.72), predation induces overcompensatory biomass responses in the consumer. This overcompensation releases the adult consumer life stage from strong intraspecific competition, resulting in larger reproductive output (SI Appendix, Fig. S8) and hence a shift in consumer stage structure (SI Appendix, Fig. S7) – see also (26). If overcompensation is necessary for predator persistence, bistability emerges in the community as predators are able to persist but not invade a stable consumer-resource system. This bistability of the community at higher temperatures occurs because consumer top down control of basal resource levels increases with warming, and the predator population then declines due to the lower basal resource levels. Below a certain equilibrium biomass density, predators cannot invade a stable consumer-resource system as the total predation pressure is not large enough to alter the stage structure of the consumer population to the predator’s favor. Specifically, persistence is possible for all temperatures below ∼33 °C (limit point, saddle node bifurcation), while invasion is only possible below ∼22 °C (branch point, transcritical bifurcation) for this specific parameter configuration (Fig. 1H). At lower temperatures, the basal resource levels, and thus predator densities, are sufficiently large for the predator to invade a consumer-resource system and hence there is no bistability.

### Varying temperature-scaling scenarios

Warming-induced alternative stable states emerge when predators feed predominantly on juveniles, both when basal resource productivity is temperature-independent (*E_R_max__* = 0) and when it declines with temperature (*E_R_max__* = −0.43) for most of the productivity values at reference temperature, 19 °C (Fig. 3; see SI Appendix, Fig. S4 for biomass densities). However, resource productivity regulates predator biomass density, which in turn determines the predator population’s ability to control the size distribution of the consumer (and, hence, ability to induce overcompensation in the consumer which is key for bistability to occur) (SI Appendix, Fig. S3). Therefore, both resource productivity and its scaling with temperature affect at which temperature transitions between different types of dynamics occur (fixed points, limits cycles or bistability) and food chain structure over temperature, such that predator persistence decreases faster with warming when the productivity declines with temperature (*E_R_max__* = −0.43) (Fig. 3B and D). The parameter combinations considered in the main analyses, based on empirical relationships, all lead to declines in predator biomass density with warming. However, a few specific scenarios can lead to increases in predator biomass density over temperature. For this to occur, the following conditions must be met: i) productivity is not decreasing with temperature, ii) resource turnover rate increases faster with temperature than consumer and predator metabolic and feeding rates, and iii) the energetic efficiency of the consumer and predator does not decline with temperature (i.e. *E_I_* ≥ *E_M,μ_*) (SI Appendix, Table S3-S4).

**Fig. 3.**
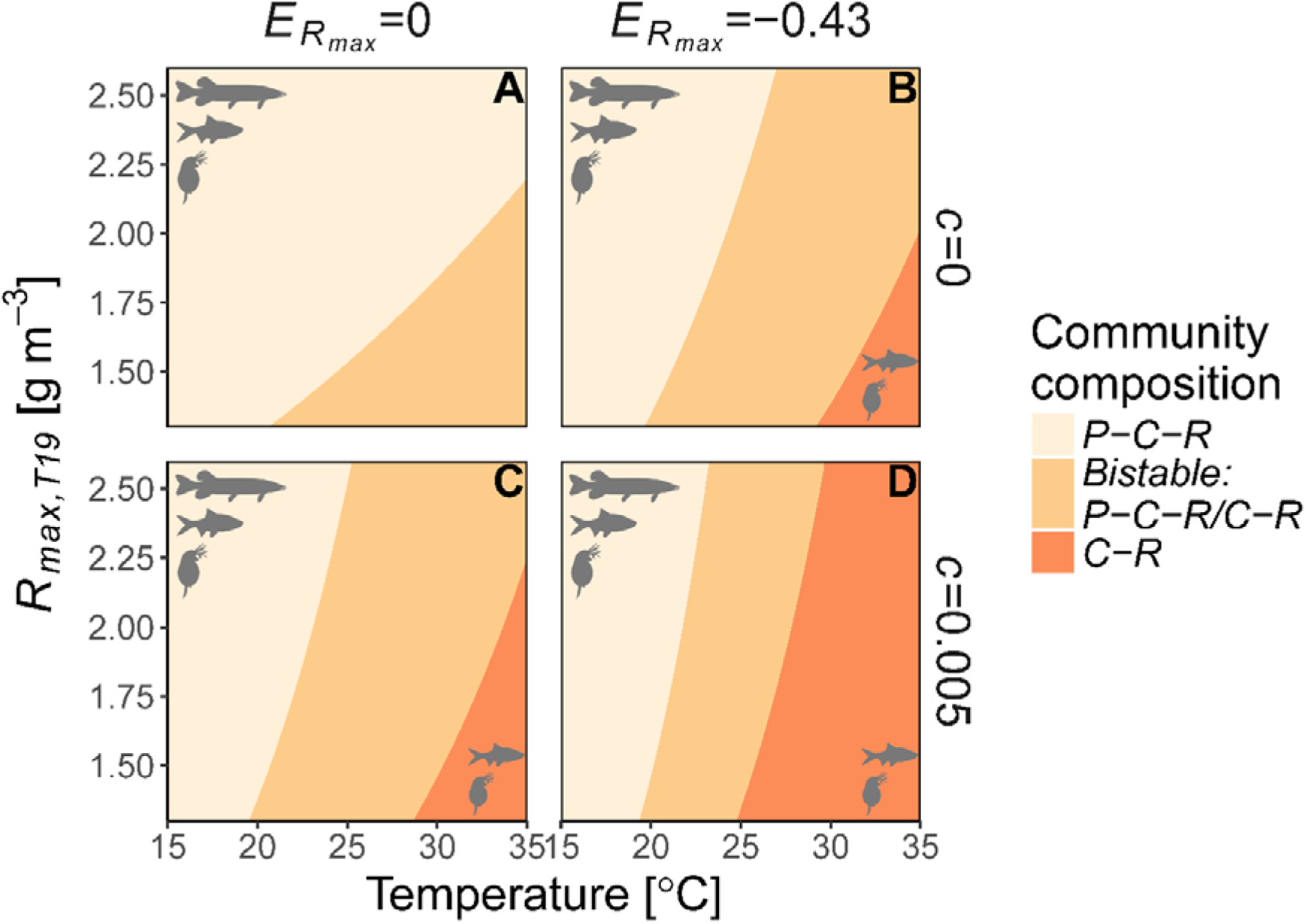
Effects of temperature on community structure depend on temperature scaling of resource (*R*) productivity and whether metabolism scales with body size and temperature independently (*c* = 0) or interactively (*c* ≠ 0) in the consumer (*C*) and predator (*P*). With warming, the tri-trophic food-chain changes from stable (off white space), to exhibiting alternative stable states (beige space; with predators either present or absent), to being reduced to two trophic levels following predator extinction (orange space). The figure shows how the species composition and dynamics of the food-chain change with temperature and resource productivity, given no (*E_R_max__* = 0) (A, C) or negative (*E_R_max__* = −0.43) (B, D) effects of temperature on resource productivity, *R*_*max,T*19_, with independent (A, B) or interactive (C, D) effects of body size and temperature on metabolism. The predator feeds exclusively on juveniles (*p* = 1), all other parameters have default values (SI Appendix, Table S1).

We model temperature-size interactions on vital rates by adding a temperature dependence (*c*) to the allometric exponent of metabolism (31) (see “*Materials and Methods*” and Eq. S1). When temperature affects the size dependence of metabolic rate (*c* > 0), the food density needed for a consumer or predator to grow (*R_crit_*) has a steeper scaling with body size compared to when assuming independent effects of temperature and body size (SI Appendix, Fig. S2). Thus, temperature-size interactions (*c* > 0) imply that energetic costs increase faster with warming in large compared to small individuals. In these scenarios (Fig. 3C-D), predator persistence is lower than in the corresponding scenario with no temperature-size interaction (Fig. 3A-B). This is because *c* > 0 reduces the growth performance more strongly in large individuals at high temperatures, which leads to a lower reproductive output of adults and a lower juvenile to adult biomass ratio (SI Appendix, Fig. S7). Therefore, temperature-size interactions negatively affect predators that feed predominantly on juveniles via two mechanisms: reductions in prey availability due to shifts in the prey size structure and increased metabolic costs. Consequently, a predator species feeding on both consumer life stages can persist at higher temperatures than one specialized on juveniles (Fig. 2), and this result is independent of the relative activation energies of feeding and metabolism (SI Appendix, Fig. S9, Table S3-S4).

The stabilizing effect of increasing temperatures that manifests when predators feed on both life stages is also shaped by the productivity of the basal resource. Specifically, the temperature at which the cyclic dynamics of the food chain switch to fixed point dynamics increases with *R*_max,T19_ (SI Appendix, Fig. S5), because the mechanism is a reversed enrichment cycle (41) and the basal resource biomass declines with warming. Factors that promote stable coexistence at high temperatures are a high and temperature-independent resource productivity (*R*_max,T19_ > 2.25) (Fig. 3A), predators feeding on both consumer life stages (Fig. 2), high energetic efficiency (SI Appendix, Table S4) and no temperature-size interactions for the scaling of metabolic rate in consumers and predators (*c* = 0, cf. Fig. 3A; SI Appendix, Fig. S2).

### Effects of warming on mean community body size

The decline in predator biomass with increased temperatures leads to a decline in the biomass-weighted mean community body size (Fig. 4). As with the abrupt predator collapse, the community size structure can also show a non-gradual abrupt decline as temperature increases, leading to alternative stable “community size states”. This decline in mean community body size occurs in all scenarios considered, and is more pronounced with interactive effects of temperature and body size on vital rates and when basal resource productivity declines with temperature (Fig. 4A-B).

**Fig. 4.**
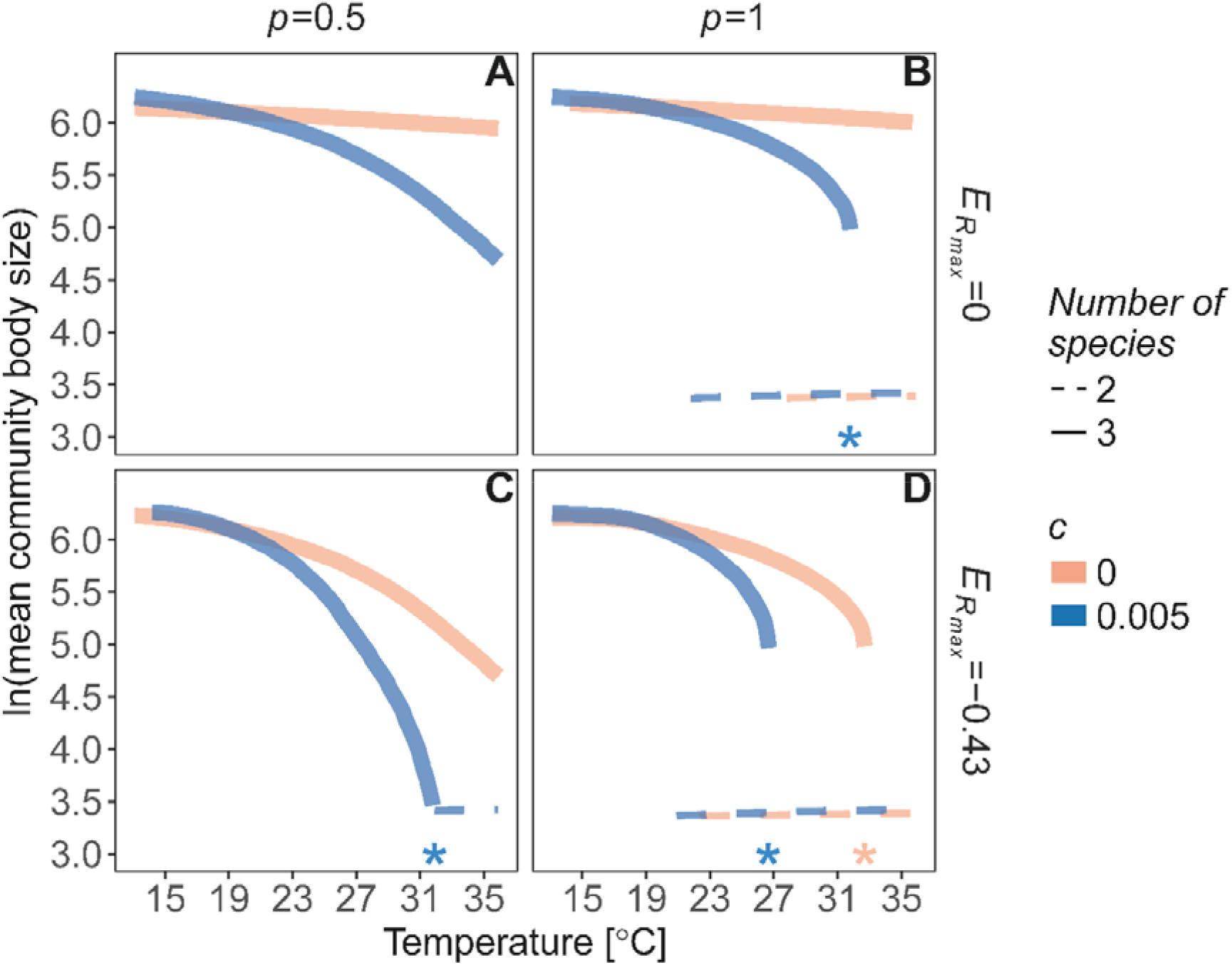
Mean body size (*S_T_*) in the community decreases with temperature, and warming can induce abrupt shifts in mean community body size. The warming effects on (*S_T_*) depend on basal productivity, ecological interactions and temperature-size interactions, as shown for food-chains with a predator species feeding with equal intensity on both consumer life stages (A, C) (*p* = 0.5) or exclusively on juveniles (B, D) *p* = 1), in a system with no temperature effects (A, B) on the basal resource productivity (*E_R_max__* = 0) or declining resource productivity with temperature (C, D) (*E_R_max__* = −0.43). Colors indicate different temperature-size scaling of metabolism, where coral lines show independent effects of body size and temperature, *c* = 0 and blue lines show positive interactive effects, *c* = 0.005. Dashed curves correspond to equilibria in which the predator has gone extinct, and mean community body size correspondingly has shifted to smaller values. Stars indicate the maximum temperature for predator persistence. Parameters have default values (SI Appendix, Table S1).

## Discussion

Here we show that food dependence of life history processes, such as maturation and reproduction, and size preference of predator feeding can explain previously unrecognized community-wide responses to warming, including alternative stable states due to emergent Allee affects, and both stabilization and destabilization of communities. These diverse community responses result from feedbacks between food, size and temperature dependence of individual performance and life history processes. We also show how warming results in declines in average community body size and reduced potential for predator persistence. These results hold across a wide range of assumptions of temperature dependencies on resource productivity, feeding and metabolism, and interactive effects of body size and temperature on metabolic rate. Much of the research on how temperature shapes population and community dynamics has focused on the role of relative temperature sensitivities of vital rates (10, 11, 13, 14). Our findings demonstrate that even under large variation in thermal sensitivities of vital rates, food- and size dependence of ecological interactions can determine the outcome of warming on community structure and dynamics. This study highlights the importance of size-based interactions and food-dependent life history processes for the energetic performance of individuals in changing climates, and how that translates to community dynamics and structure.

The general prediction of reduced predator persistence with increasing temperatures corroborates earlier studies (8, 10). Moreover, the parameter space that cause predators to decline, i.e. a combination of negative temperature dependence of resource productivity, slower resource growth rates relative to consumer and predator feeding and metabolic rates, and reduced energetic efficiency of consumers and predators, have also been found in both terrestrial and aquatic systems (5, 7–9, 12, 42–44). Our novel finding is that predator declines with increasing temperatures are not always gradual, but can be sudden and collapses can occur at temperatures much lower than those that would cause starvation. This happens when warming induces bistability due to Allee effects (26), which emerge from feedbacks between individual performance and population size structure, given food-dependent growth – all of which are ubiquitous in natural food webs. Therefore, our results suggest that warming may expose predators to an additional, previously overlooked risk of sudden collapse and/or impaired recovery potential in warmer environments. This occurs as warming can cause community bistability, in which case the predator population can collapse before energetic starvation, depending on the initial productivity and temperature of the system. This increased risk could manifest itself in systems where strong interactions among and within species shape the community size structure (the prerequisites for emergent Allee effects) as has been suggested for populations of Atlantic cod, brown trout and Arctic charr (22, 26–28). The model predicts non-gradual declines in biomass when predators feed mainly on smaller consumers. Such feeding behavior has empirical support for e.g. Atlantic salmon (*Salmo salar* L.) (45) and Atlantic cod (27, 28). Thus, the model used here generally predicts lower persistence of predators in warmer environments, and importantly, that both predator densities and mean community body size do not necessarily decline gradually with temperature but can exhibit sudden collapses.

Recent studies have aimed to reconcile the diverse effects warming can have on e.g. community stability by deriving general principles based on the relative temperature-sensitivities of resource energetic efficiency (i.e. biomass gains vs losses) and productivity (10–12). However, we do not know if these predictions apply to size-structured populations with individuals exhibiting food-dependent growth, development and reproduction. This is a key question to address, as both size variation within species and food dependence of development and reproduction are widespread in nature, and often govern ecological dynamics (15). We account for food dependence of growth, development and reproduction mechanistically, such that these processes depend on both size-dependent interactions within and among species, and on direct physiological effects of warming. The model analyses suggest that for a predator feeding on both consumer life stages, warming shifts the community from exhibiting limit cycles to stable point dynamics via an inverse paradox of enrichment mechanism, as reported also in previous studies (40, 46). In contrast to models that do not account for size-variation within populations (11), we show that the qualitative shifts in community structure and stability with increased temperatures (in terms of periodic vs fixed point equilibrium solutions, or presence of alternative stable states), are not primarily driven by the relative temperature dependence of resource productivity, feeding and metabolism. Instead, we found that the effect of body size on competitive interactions within the consumer species and the effect of predation on consumer biomass indeed can determine the effects of warming on community structure and stability.

As the effects of temperature on performance tend to vary with body size or life stage, a single activation energy parameter cannot describe the entire temperature dependence of a given vital rate (31, 47–49). Such interactive effects of temperature and body size are reflected in empirical patterns such as the temperature size rule (TSR) (increased juvenile growth- or developmental rates but smaller adult body size in warmer environments) (34), which is especially strong in aquatic environments (29, 30). Still, the implications of such temperature-size interactions for population and community dynamics are poorly understood (31). Recent studies suggest that when the average body size of species declines with warming, stability, in terms of return times after perturbations, generally increases (39). Persistence of species in a food chain generally increases when warming causes reductions in size, though this depends on the trophic level at which the reductions occur (40). In our study we find the opposite. Predator persistence is always lower with temperature-size interactions compared to independent temperature and body size effects. A key difference in our approach to previous studies of temperature and body size interactions (39, 40), in addition to accounting for stage structure within species, is that we do not assume temperature-effects on body size based on empirical temperature-size patterns. Instead, we assess how the energetic performance of an individual of a specific size changes with temperature, based on the scaling of individual-level rates, such as feeding and metabolism, and species interactions. Thus, any changes in average body size within species and of the community as a whole emerge in the model as a result of the temperature dependence of physiological rates and how temperature effects are mediated by ecological interactions.

Predators and consumers coexist when basal productivity is high, irrespective of the predator feeding preference (in stable fixed point or cyclic dynamics, respectively). Information about productivity is thus important for qualitative and quantitative predictions about the effects of warming. Given its effect on community composition, we choose values that allowed for stable coexistence at our reference temperature, given all other sets of parameters (22). There are, however, other important aspects of productivity and temperature than their numerical values. For instance, a recent study showed that responses of carrying capacity of algae to increased temperature can be increasing, decreasing or hump-shaped, if viewed as a dynamic property rather than a parameter (11). This suggests that resource densities may in fact show more complex response to temperature than acknowledged here (e.g. Fig. 3; SI Appendix, Fig. S3). We made simplifying assumptions about the resource dynamics to focus on the mechanistic feedbacks between individual-level energetics and ecological interactions in the food chain. In addition, by using the Arrhenius equation (i.e. exponential temperature dependencies) we have not accounted for hump-shaped relationships between temperature and biological rates (50, 51), which can have major effects on the stability and persistence responses to warming (11). Thereby, we likely overestimate the specific temperatures beyond which predators cannot persist. However, we do not aim to quantitatively represent a specific system but identify qualitative community responses to warming and their underlying mechanisms. Importantly, we show that in most temperature scaling scenarios considered (Fig. 3), warming-induced alternative stable states emerge in juvenile specialized predators regardless of how productivity scales with temperature.

Within-species phenotypic variation is increasingly recognized as an important driver of ecological dynamics (15, 52–55), and there is vast empirical evidence on size-dependent responses to warming (16, 17, 34). Accordingly, warming should influence the outcome of size-dependent interactions. However, the ecological implications of within species variation in the context of responses to global warming have been largely overlooked (but see (31, 38)). Our results demonstrate that approaches based on species-averaged traits (such as mean body size) cannot accurately represent the full range of dynamics and shifts in community size structure that warming is causing globally. We show that even simple stage structure within a population can result in unexpected community-level responses to rising temperatures, including alternative stable food web states and potential collapses of predators due to emergent Allee effects. Thus, feedbacks between food dependence of life history processes and population size structure, both ubiquitous in natural food webs, can alter the effects of temperature on food web stability and predator persistence.

## Materials and Methods

### Modeling framework

To study the effects of warming on coexistence and stability of a tri-trophic food chain taking within-species stage structure and size- and food-dependent processes into account, we used a stage-structured biomass model (Eqns. 1–4) (56). Stage-structured biomass models are derived from – and under equilibrium conditions exactly represent – size-structured population models with a continuous size distribution (57). Following bioenergetics and mass conservation principles (58), biomass production (used for growth, development and reproduction) is food-dependent. Consequently, growth performance is mediated by ecological interactions within and between species via exploitation of shared resources.

The model in this paper is an extension of the temperature-dependent consumer-resource model used in (31), empirically parameterized to represent a stage-structured consumer zooplanktivorous fish (common roach, *Rutilus rutilus* L.) and its zooplankton prey (*Daphnia sp*.), here extended with a size-selective predator (pike, *Esox Lucius* L.) feeding on the consumer. The life stages considered in the consumer are juveniles and adults, as maturation and reproduction are two of the most fundamental life history processes in animals (15). Juvenile and adult consumers and predators are characterized by a representative body size (∼4 g, ∼30 g and ∼640 g, respectively; see SI Appendix), which are used to calculate their average mass-specific rates of metabolism, maximum ingestion, attack and background mortality. We also account for interactive effects of body size and temperature on metabolic rate to approximate the increasing costs of being large in a warmer environment, see *“Size- and temperature dependence of vital rates”*, below for details. Independent of temperature scaling, asymmetrical competition between life stages in the consumer population arises from differences in body size and thus energetic performance (i.e. energetic gains minus losses from metabolism and mortality). The state variables are biomass densities [g m^-3^] of a basal resource, juvenile and adult consumers feeding on the resource and a predator feeding with varied size preference on consumers (referred to as *R, J, A* and *P*, respectively) (Eqns. 1–4): 

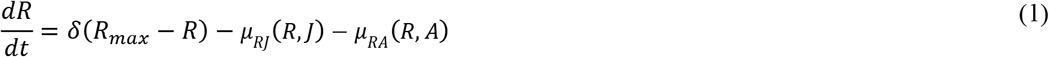

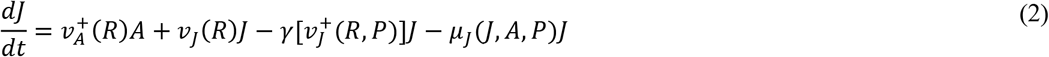

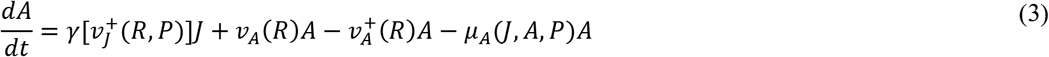

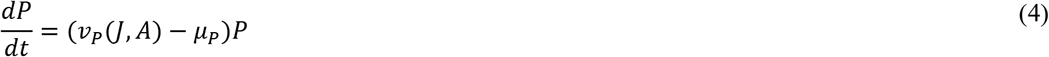

Most terms in the system of ordinary differential equations are species- and mass-specific functions of body size, temperature and the basal resource. These are described in the following paragraphs and in Table 1. For parameters and derivation of the allometric relationships within the functions, we refer to SI Appendix, Table S1.

We assume that the basal resource (*R*) grows according to semi-chemostat dynamics (59), with temperature-dependent turnover rate (*δ*) and maximum density (*R*_max,T19_). Juvenile biomass increases with adult reproduction (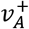). The ^+^-superscript refers to positive values of biomass production such that reproduction and maturation only occurs when biomass is produced, which ensures that starvation (i.e. when *v_J,A_* < 0) in one life stage does not reduce biomass of the other life stage. However, since we analyze equilibrium dynamics, starvation is not possible without consumer extinction, which we did not encounter in any of our modeled scenarios. Juvenile biomass is lost through maturation (*γ*) into the adult stage and mortality, *μ_J_* (sum of background and predation mortality). Adult biomass is gained through maturation (*γ*) and lost through reproduction (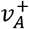) and background and predation mortality (*μ*_*A*_). We assume that juveniles invest all energy into growth and development, whereas adults spend all their energy on reproduction and hence do not grow in size (56). The predator population is unstructured as we are primarily interested in how warming responses are influenced by the relation between consumer size structure and mortality imposed by the predator. This does not necessarily rely on structure in the predator, but rather on the interplay between size-specific predation mortality and stage-structured dynamics in the consumer (26). Hence, the biomass dynamics of the predator are described by its temperature-dependent net energy production (*v*_*p*_) and losses due to background mortality *μ*_*P*_.

The net biomass production of consumers and predators (*v_J,A,P_*) is the difference between ingested energy, scaled with assimilation efficiency *σ*_*z,p*_ (Table 1; SI Appendix, Table S1), and metabolic costs (*M_J,A,P_*). Ingestion follows a Holling type II functional response (60) for consumers and predators, with size- and temperature-dependent functions describing maximum ingestion (*I_max,J,A,P_*) and attack rate (*α_J,A,P_*) (Table 1; SI Appendix). We vary the feeding preference of predators by scaling their encounter rate of juveniles with parameter *p* and of adults with 1 − *p* (i.e. *p* = 1 means juvenile selective predator, *p* = 0.5 no preference and *p* = 0 means adult selective predator).

**Table 1.**
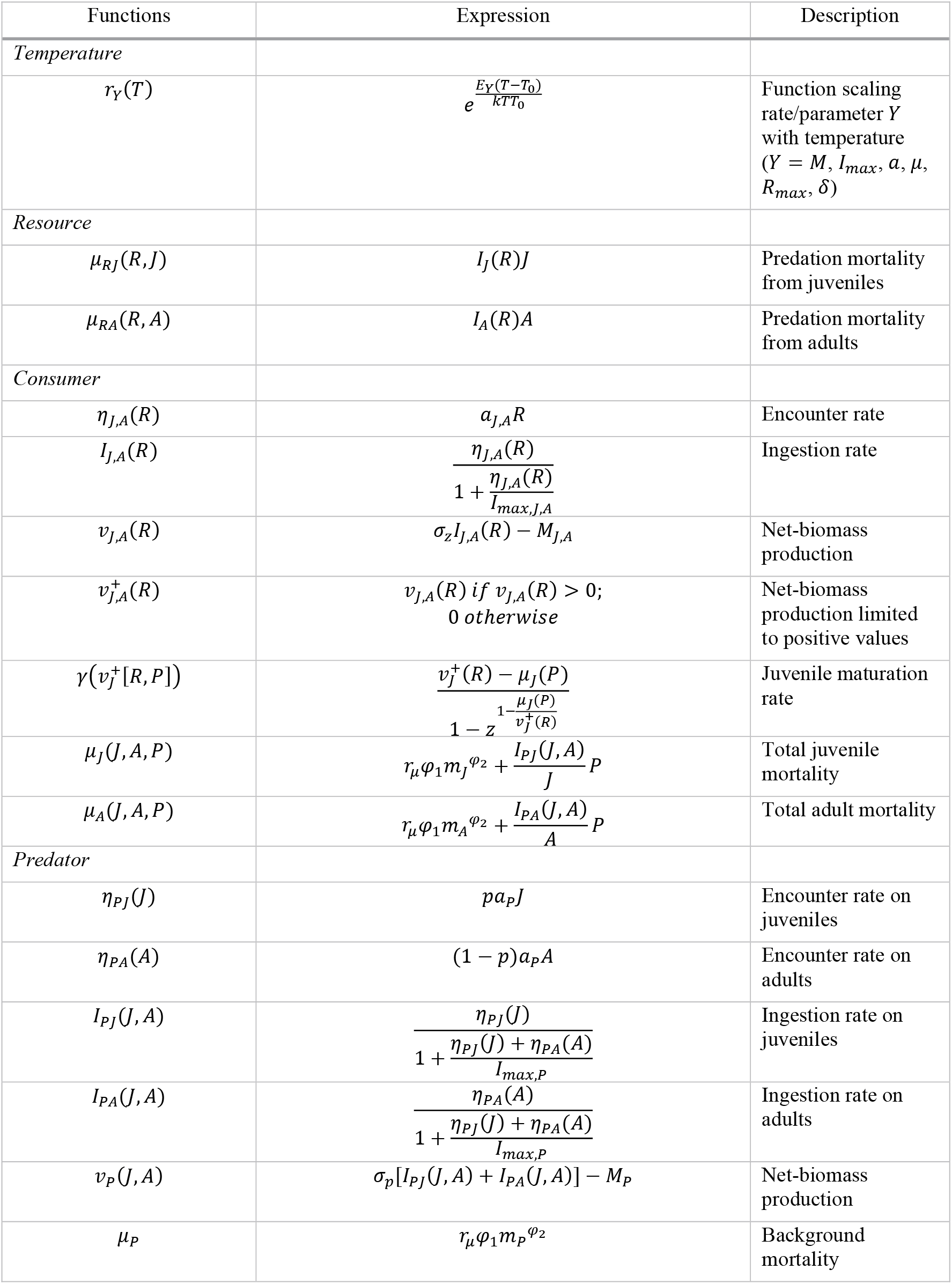
Model functions. Note that dependencies are only expressed for state variables, but all functions relate to individual-level rates that depend both on body size and temperature (see SI Appendix, Table S1).

### Size- and temperature dependence of vital rates

All individual-level rates are functions of both body size and temperature (Table 1; see also SI Appendix for derivation of functions and Table S1 for their numerical values at 19 °C). Temperature dependence is acquired by multiplying allometric functions of mass-specific rates (at reference temperature, 19 °C) with a Boltzmann-Arrhenius function, scaled to equal 1 at 19 °C (31). Allometric functions are of the form *aM^b^*, with species- and rate-specific constants and exponents, *a* and *b*, respectively. Attack rates are derived from more complex, hump-shaped functions that depend on both predator and prey body sizes (59, 61, 62) (SI Appendix, Table S1). The Boltzmann-Arrhenius function is given by 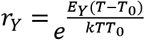, where *T* [K] is the temperature, *T*_0_ [K] is the reference temperature, *k* [eVK^-1^] is Boltzmann’s constant and *E_Y_* [eV] is the activation energy of rate or parameter *Y* (63). These temperature-dependent rates and parameters (*Y*) include: metabolism (*M_J,A,P_*), functional response parameters attack rate (*a_J,A,P_*) and maximum ingestion rate (*I_max,J,A,P_*), basal resource turnover rate (*δ*), maximum resource density (*R*_max,T19_) and background mortality (*μ_J,A,P_*).

When varying the productivity of the basal resource (*R*_max,T19_) and its scaling with temperature through its activation energy, *E_R_max__*, two contrasting scenarios were considereds: (i) no effect of temperature on *R*_max,T19_ (*E_R_max__* = 0), or (ii) *R*_max,T19_ declining with temperature at the same rate as turnover rate increases (*E_R_max__* = −*E*_*δ*_ = −0.43 [eV]) (SI Appendix). This assumption is based on mass conservation and metabolic scaling principles (13, 39, 64). Given the large variation in activation energies of feeding rates in the literature, in particular within species (50, 51), we varied the activation energy of functional response parameters (*E_I_*) by scaling it with a factor ranging between 0.5 and 1.5 relative to the activation energy of metabolism (*E*_*M*_) (SI Appendix, Table S1, S4; Fig. S9-S10). This results in *E_I_*-values between 0.297 and 0.891, which are in the range of estimates reported in the literature (43, 50, 51).

Thus far, the formulation follows closely that of the metabolic theory of ecology (1), in that temperature effects are exponential and independent of body size. However, we also relax the assumption of independent effects of body size and temperature. This is done by adding a linear temperature dependence (*c*) to the exponent of metabolic rate, such that 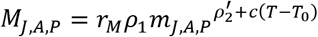, where *r_M_* is the temperature scaling factor for metabolism, *ρ*_1_ is the normalization constant, *m_J,A,P_* is the mass of juveniles, adults or predators and 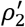 is the allometric exponent at *T*_0_ (292 °K, i.e. 19 °C), and *c* determines whether metabolism increases faster with temperature for large relative to small individuals (*c* > 0), or vice versa (*c* < 0) (SI Appendix, Fig. S2). This linear form of temperature-size interaction has been shown within species (31, 49). We assume that both consumers and predators have the same *c* by default (but see SI Appendix, Fig. S9), but acknowledge that it can vary considerably between species (31, 49). A steeper size scaling of metabolism with body size at higher temperatures (*c* > 0), all else equal, leads to a steeper scaling of the critical resource density needed to meet metabolic demands (*R_crit_*) with body size (SI Appendix, Fig. S2). Therefore, an interactive effect of body size and temperature on metabolism (*c* > 0) results in a reduced growth performance for larger individuals in warmer environments, as is often observed in aquatic systems (29, 30). *R_crit_* is given by 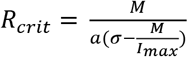, where *M* is metabolic rate, *σ* is the assimilation efficiency, *a* is attack rate and *I_max_* is maximum ingestion rate (65). Note also that steeper scaling of critical resource density (*R_crit_*) with body size can arise with *c* ≤ 0, as long as the size-scaling exponent of feeding rate decreases more rapidly with temperature than that of the metabolic exponent (SI Appendix, Fig. S2).

Importantly, we do not make assumptions about temperature effects on integrated traits such as body size, but instead model interactive temperature-size effects mechanistically on individual-level rates from which body growth results. Thus, effects of warming stem from direct effects on individual-level processes that are mediated by ecological interactions within and between species via exploitation of shared resources.

### Analysis

We analyzed equilibrium biomass densities and bifurcations by performing equilibrium continuations, which allows for also tracing unstable equilibria, using the MATLAB (66) package MATCONT GUI (67). In using numerical techniques, the specific results presented here (e.g. temperatures at bifurcations) should be viewed in relation to the parameter setup and not as quantitative predictions for a particular system. Therefore, we test the robustness of our results in various sensitivity analyses, including a wide variety of temperature dependencies. Minimum and maximum of limit cycles were acquired by running time integrations until equilibrium for selected parameters and with continuation analysis, and the stability of limit cycles was tested using Floquet theory (68). We also calculated biomass-weighted mean body size (*S_T_*) of the community for stable equilibria at temperature *T* (averaged for each *T*) as 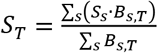, where *S_s_* is the representative body size of juvenile consumers, adult consumers or predators (*s* = *J,A,P*) and *B_s,T_* is their corresponding equilibrium biomass density at temperature *T*. Model files, instructions for viewing and implementing the model in MATCONT, and R-scripts to reproduce the main figures with simulated data have been deposited on https://github.com/maxlindmark/Temperature_Allee. R version 3.4.2 (69) was used to create figures.

## Acknowledgements

We thank Sebastian Diehl, Wojciech Uszko and Viktor Thunell for inspiring and helpful discussions, André De Roos for assistance in implementing the maturation function and phylopic (http://phylopic.org/) for the pike silhouette, available for reuse under the Public Domain Mark 1.0 license (the other silhouettes were drawn by ML). This work was supported by grants from the Swedish Research Council (no. 2015-03752 to AG), and the Swedish Research Council FORMAS (no. 217-2013-1315 to AG).

## Author contributions

AG conceived the study. All authors contributed to the formulation of research questions and study design. ML performed modelling analyses and wrote the first draft. All authors contributed to the revision of the manuscript.

## Cited Literature

1. Brown JH, Gillooly JF, Allen AP, Savage VM, West GB (2004) Toward a metabolic theory of ecology. Ecology 85(7):1771–1789.

2. Savage VM, Gillooly JF, Brown JH, West GB, Charnov EL (2004) Effects of body size and temperature on population growth. Am Nat 163(3):429–441.

3. Peters RH (1983) The ecological implications of body size (Cambridge University Press).

4. Blanchard JL, et al. (2012) Potential consequences of climate change for primary production and fish production in large marine ecosystems. Philos Trans R Soc Lond B Biol Sci 367(1605):2979–2989.

5. O’Gorman EJ, et al. (2017) Unexpected changes in community size structure in a natural warming experiment. Nat Clim Change 7(9):659–663.

6. Yvon-Durocher G, Montoya JM, Trimmer M, Woodward GUY (2011) Warming alters the size spectrum and shifts the distribution of biomass in freshwater ecosystems. Glob Change Biol 17(4):1681–1694.

7. O’Connor MI (2009) Warming strenghtens an herbivore-plant interaction. Ecology 90(2):388–398.

8. Rall BC, Vucic-Pestic O, Ehnes RB, Emmerson M, Brose U (2010) Temperature, predator-prey interaction strength and population stability. Glob Change Biol 16(8):2145–2157.

9. Vucic-Pestic O, Ehnes RB, Rall BC, Brose U (2011) Warming up the system: higher predator feeding rates but lower energetic efficiencies. Glob Change Biol 17(3):1301–1310.

10. Fussmann KE, Schwarzmüller F, Brose U, Jousset A, Rall BC (2014) Ecological stability in response to warming. Nat Clim Change 4(3):206–210.

11. Uszko W, Diehl S, Englund G, Amarasekare P (2017) Effects of warming on predator-prey interactions: a resource-based approach and a theoretical synthesis. Ecol Lett 20(4):513–523.

12. Vasseur DA, McCann KS (2005) A mechanistic approach for modelling temperature-dependent consumer-resource dynamics. Am Nat 166(2):184–198.

13. Gilbert B, et al. (2014) A bioenergetic framework for the temperature dependence of trophic interactions. Ecol Lett 17(8):902–914.

14. O’Connor MI, Gilbert B, Brown CJ (2011) Theoretical predictions for how temperature affects the dynamics of interacting herbivores and plants. Am Nat 178(5):626–638.

15. Persson L, De Roos AM (2013) Symmetry breaking in ecological systems through different energy efficiencies of juveniles and adults. Ecology 94(7):1487–1498.

16. Angilletta MJ, Dunham AE (2003) The temperature-size rule in ectotherms: simple evolutionary explanations may not be general. Am Nat 162(332–342).

17. Pörtner HO, Farrell AP (2008) Physiology and climate change. Science 322(5902):690–692.

18. Ohlberger J (2013) Climate warming and ectotherm body size – from individual physiology to community ecology. Funct Ecol 27(4):991–1001.

19. De Roos AM, et al. (2007) Food ‐dependent growth leads to overcompensation in stage ‐specific biomass when mortality increases: the influence of maturation versus reproduction regulation. Am Nat 170(3):E59–E76.

20. Huss M, Nilsson KA (2011) Experimental evidence for emergent facilitation: promoting the existence of an invertebrate predator by killing its prey. J Anim Ecol 80(3):615–621.

21. Ohlberger J, et al. (2011) Stage-specific biomass overcompensation by juveniles in response to increased adult mortality in a wild fish population. Ecology 92(12):2175–2182.

22. Persson L, et al. (2007) Culling prey promotes predator recovery – alternative states in a whole-lake experiment. Science 316(5832):1743–1746.

23. Schröder A, Persson L, de Roos AM (2009) Culling experiments demonstrate size-class specific biomass increases with mortality. Proc Natl Acad Sci USA 106(8):2671–2676.

24. Schröder A, van Leeuwen A, Cameron TC (2014) When less is more: positive population-level effects of mortality. Trends Ecol Evol 29(11):614–624.

25. De Roos AM, Persson L, Thieme HR (2003) Emergent Allee effects in top predators feeding on structured prey populations. Proc R Soc Lond B Biol Sci 270(1515):611–618.

26. De Roos AM, Persson L (2002) Size-dependent life-history traits promote catastrophic collapses of top predators. Proc Natl Acad Sci USA 99(20):12907–12912.

27. Gårdmark A, et al. (2015) Regime shifts in exploited marine food webs: detecting mechanisms underlying alternative stable states using size-structured community dynamics theory. Philos Trans R Soc Lond B Biol Sci 370(1659):20130262.

28. van Leeuwen A, De Roos AM, Persson L (2008) How cod shapes its world. J Sea Res 60(1–2):89–104.

29. Forster J, Hirst AG, Atkinson D (2012) Warming-induced reductions in body size are greater in aquatic than terrestrial species. 109(47):19310–19314.

30. Horne CR, Hirst AG, Atkinson D (2015) Temperature-size responses match latitudinal-size clines in arthropods, revealing critical differences between aquatic and terrestrial species. Ecol Lett 18(4):327–335.

31. Lindmark M, Huss M, Ohlberger J, Gårdmark A (2018) Temperature-dependent body size effects determine population responses to climate warming. Ecol Lett 21(2):181–189.

32. Malerba ME, White CR, Marshall DJ (2018) Eco-energetic consequences of evolutionary shifts in body size. Ecol Lett 21(1):54–62.

33. Reuman DC, Holt RD, Yvon-Durocher G (2014) A metabolic perspective on competition and body size reductions with warming. J Anim Ecol 83(1):59–69.

34. Atkinson D (1994) Temperature and Organism Size—A Biological Law for Ectotherms? Adv Ecol Res 25:1–58.

35. Kingsolver JG, Huey RB (2008) Size, temperature, and fitness: three rules. Evol Ecol Res 10:251–268.

36. Björnsson B, Steinarsson A (2002) The food-unlimited growth rate of Atlantic cod (Gadus morhua). Can J Fish Aquat Sci 59(3):494–502.

37. Karås P, Thoresson G (1992) An application of a bioenergetics model to Eurasian perch (Perca fluviatilis L.). J Fish Biol 41:217–230.

38. Ohlberger J, Edeline E, Vollestad LA, Stenseth NC, Claessen D (2011) Temperature-driven regime shifts in the dynamics of size-structured populations. Am Nat 177(2):211–223.

39. Osmond MM, et al. (2017) Warming-Induced Changes to Body Size Stabilize Consumer-Resource Dynamics. Am Nat 189(6):718–725.

40. Sentis A, Binzer A, Boukal DS (2017) Temperature-size responses alter food chain persistence across environmental gradients. Ecol Lett 20(7):852–862.

41. Rosenzweig ML (1971) Paradox of Enrichment: Destabilization of Exploitation Ecosystems in Ecological Time. Science 171(3969):385–387.

42. Downs CJ, Hayes JP, Tracy CR (2008) Scaling metabolic rate with body mass and inverse body temperature: A test of the Arrhenius fractal supply model. Funct Ecol 22(2):239–244.

43. Rall BC, et al. (2012) Universal temperature and body-mass scaling of feeding rates. Philos Trans R Soc Lond B Biol Sci 367(1605):2923–2934.

44. Segura AM, Sarthou F, Kruk C (2018) Morphology-based differences in the thermal response of freshwater phytoplankton. Biol Lett 14(5):20170790.

45. Jacobson P, Gårdmark A, Östergren J, Casini M, Huss M (2018) Size-dependent prey availability affects diet and performance of predatory fish at sea: a case study of Atlantic salmon. Ecosphere 9(1):e02081.

46. Binzer A, Guill C, Brose U, Rall BC (2012) The dynamics of food chains under climate change and nutrient enrichment. Philos Trans R Soc Lond B Biol Sci 367(1605):2935–2944.

47. Xie X, Sun R (1990) The Bioenergetics of the Southern Catfish (Silurus meridionalis Chen). I. Resting Metabolic Rate as a Function of Body Weight and Temperature. Physiol Zool 63(6):1181–1195.

48. Beamish FWH (1964) Respiration of fishes with special emphasis on standard oxygen consumption: II. Influence of weight and temperature on respiration of several species. Can J Zool Can Zool 42(2):177–188.

49. Ohlberger J, Mehner T, Staaks G, Hölker F (2012) Intraspecific temperature dependence of the scaling of metabolic rate with body mass in fishes and its ecological implications. Oikos 121(2):245–251.

50. Dell AI, Pawar S, Savage VM (2011) Systematic variation in the temperature dependence of physiological and ecological traits. Proc Natl Acad Sci USA 108(26):10591–10596.

51. Englund G, Öhlund G, Hein CL, Diehl S (2011) Temperature dependence of the functional response. Ecol Lett 14(9):914–921.

52. Bolnick DI, et al. (2011) Why intraspecific trait variation matters in community ecology. Trends Ecol Evol 26(4):183–192.

53. Des Roches S, et al. (2018) The ecological importance of intraspecific variation. Nat Ecol Evol 2(1):57–64.

54. Miller TEX, Rudolf VHW (2011) Thinking inside the box: Community-level consequences of stage-structured populations. Trends Ecol Evol 26(9):457–466.

55. Ryabov AB, De Roos AM, Meyer B, Kawaguchi S, Blasius B (2017) Competition-induced starvation drives large-scale population cycles in Antarctic krill. Nat Ecol Evol 1:0177.

56. De Roos AM, et al. (2008) Simplifying a physiologically structured population model to a stage-structured biomass model. Theor Popul Biol 73(1):47–62.

57. Metz JAJ, Diekmann O (1986) The dynamics of physiologically structured populations (Springer-Verlag, Heidelberg, Germany).

58. Yodzis P, Innes S (1992) Body size and consumer-resource dynamics. Am Nat 139(6):1151–1175.

59. Persson L, Leonardsson K, De Roos AM, Gyllenberg M, Christensen B (1998) Ontogenetic scaling of foraging rates and the dynamics of a size-structured consumer-resource model. Theor Popul Biol 54:270–293.

60. Holling CS (1959) Some characteristics of simple types of predation and parasitism. Can Entomol 91(07):385–398.

61. Hjelm J, Persson L (2001) Size-dependent attack rate and handling capacity: inter-cohort competition in a zooplanktivorous fish. Oikos 95:520–532.

62. Claessen D, De Roos AM, Persson L (2000) Dwarfs and Giants: Cannibalism and Competition in Size ‐Structured Populations. Am Nat 155(2):219–237.

63. Gillooly JF, Brown JH, West GB, Savage VM, Charnov EL (2001) Effects of size and temperature on metabolic rate. Science (293.5538):2248–2251.

64. Bernhardt JR, Sunday JM, O’Connor MI (2018) Metabolic theory and the temperature size rule explain the temperature dependence of population carrying capacity. Am Nat In press. doi:https://dx.doi.org/10.1086/700114.

65. Byström P, Andersson J (2005) Size ‐dependent foraging capacities and intercohort competition in an ontogenetic omnivore (Arctic char). Oikos 110(3):523–536.

66. MATLAB (2014) version 8.4.0.150421 (2014b). Natick, Ma.

67. Dhooge A, Govaerts W, Kuznetsov YA, Meijer HGE, Sautois B (2008) New features of the software MatCont for bifurcation analysis of dynamical systems. Math Comput Model Dyn Syst 14(2):147–175.

68. Klausmeier CA (2008) Floquet theory: a useful tool for understanding nonequilibrium dynamics. Theor Ecol 1(3):153–161.

69. R Core Team (2018) R: A Language and Environment for Statistical Computing. R Foundation for Statistical Computing (Vienna, Austria) Available at: https://www.R-project.org/.

